# Motor and cognitive sequence tasks exhibit different ramping patterns in parietal and prefrontal cortices

**DOI:** 10.1101/2024.10.09.617499

**Authors:** Hannah Doyle, Rhys Yewbrey, Katja Kornysheva, Theresa M. Desrochers

## Abstract

Humans complete different types of sequences as a part of everyday life. These sequences can be divided into two important categories: those that are abstract, in which the steps unfold according to a rule at super-second to minute time scale, and those that are motor, defined solely by individual movements and their order which unfold at the sub-second to second timescale. For example, the sequence of making spaghetti consists of abstract tasks (preparing the sauce and cooking the noodles) and nested motor actions (stir pasta water). Previous work shows neural activity increases (ramps) in the rostrolateral prefrontal (RLPFC) during abstract sequence execution (Desrochers et al., 2015, 2019). During motor sequence production, activity occurs in regions of the prefrontal cortex (Yewbrey et al., 2023). However, it remains unknown if ramping is a signature of motor sequence production as well or solely an attribute of abstract sequence monitoring and execution. We tested the hypothesis that significant ramping activity occurs during motor sequence production in the RLPFC. Contrary to our hypothesis, we did not observe significant ramping activity in the RLPFC during motor sequence production, but we found significant ramping activity in bilateral inferior parietal cortex, in regions distinct from those observed during an abstract sequence task. Our results suggest different prefrontal-parietal circuitry may underlie abstract versus motor sequence execution.

## Introduction

Achieving goals in everyday life requires completing sequences of tasks. For example, baking a cake requires following instructions provided by a recipe. The steps of a recipe can be broken down into different sequence types. The abstract sequence details ordered tasks without specific motor actions, such as combining the dry ingredients and preparing the pan. Each of these tasks is comprised of motor sequences of actions, like scooping out the flour and brushing the pan with butter. Thus, sequences can be considered to contain both abstract and motor components. Abstract sequences are defined by a rule that governs the order but not motor identity of steps (Desrochers et al., 2022), while motor sequences, which can be nested into the abstract sequence, are defined by a specific series of actions in a certain order (Hikosaka et al., 2002; Rosenbaum et al., 2007; Yewbrey et al., 2023). Neurally, the rostrolateral prefrontal cortex (RLPFC, also referred to as lateral anterior prefrontal cortex) has been shown to be necessary for abstract sequences (Desrochers et al., 2015), but the role of this region in motor sequences remains unclear. Because motor and abstract sequence execution often occur together, a common sequence tracking process in the RLPFC may underlie them. The present study tested this possibility by examining neural dynamics in the PFC during motor sequence production compared to abstract sequences.

Motor sequences such as stirring (**Figure 1A**), are supported primarily by activity in motor, premotor and parietal cortices. The premotor, supplementary motor, and parietal cortices contain information about chunks and position rank in motor sequences (Tanji and Shima, 1994; Yokoi and Diedrichsen, 2019; Russo et al., 2020). The motor cortex is associated with the execution of single movements in sequences (Kornysheva and Diedrichsen, 2014; Yokoi et al., 2018; Zimnik and Churchland, 2021; Ariani et al., 2022; Yewbrey et al., 2023). Although some electrophysiological studies in non-human primates (Averbeck et al., 2002) and neuroimaging studies in humans (Kornysheva and Diedrichsen, 2014; Yokoi et al., 2018; Yokoi and Diedrichsen, 2019) describe prefrontal cortex activity related to motor sequence production, most literature focuses on activity in motor related and parietal cortices. Literature thus far shows motor and related cortical regions support motor sequence execution, and the potential contribution of the RLPFC in this process remains unclear.

**Figure 1.**
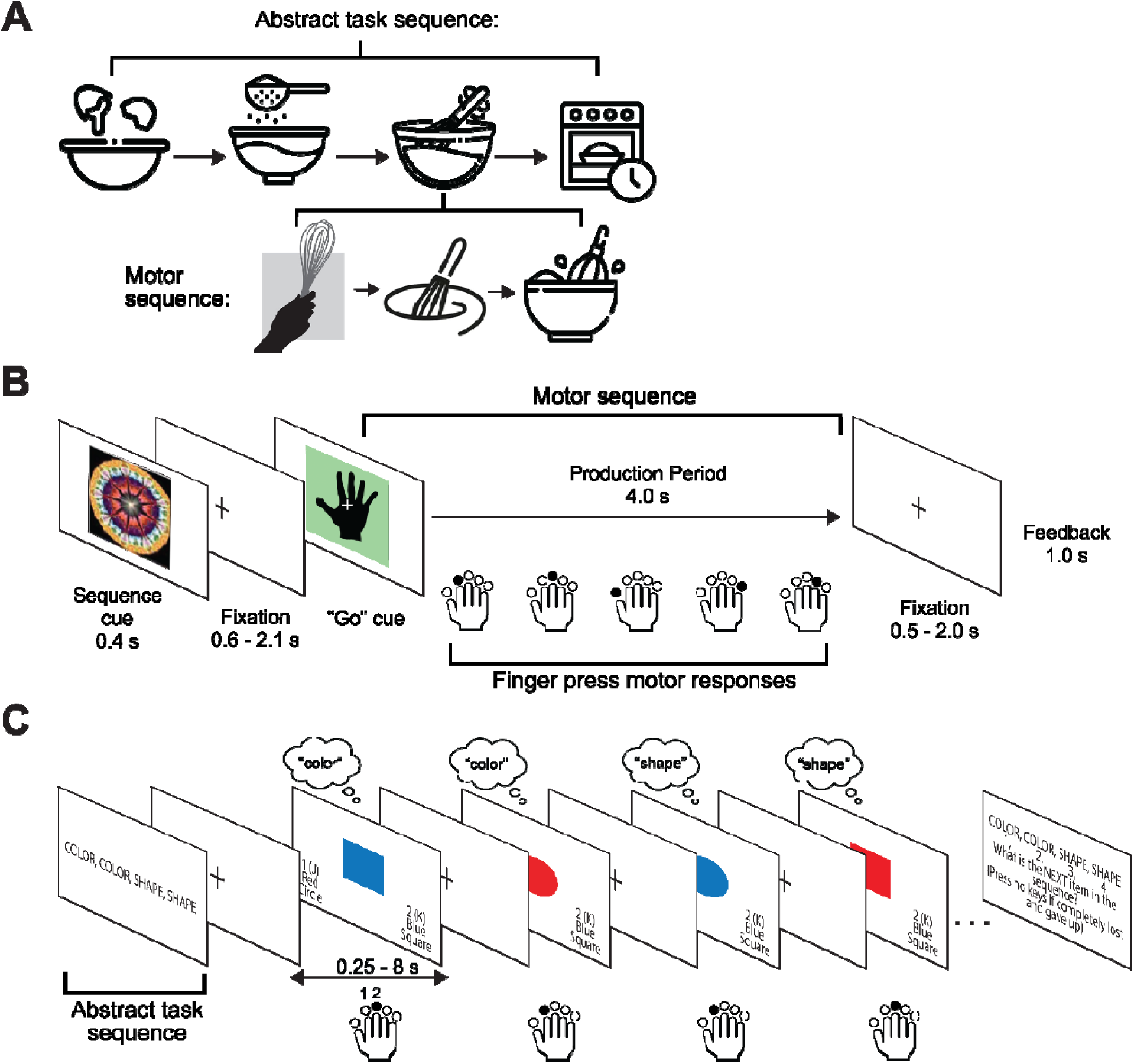
Abstract and motor sequence schematic and behavioral tasks. **A.** Abstract sequences contain tasks that do not rely on specific motor actions to complete (e.g., tasks of adding eggs, sifting flour, combining ingredients, and putting them in the oven when baking a recipe). Motor sequences contain tasks that require specific actions to achieve the goal (e.g., to whisk together all the ingredients one might grab the whisk, make a stirring motion, and tap the whisk against the bowl) **B.** The motor sequence paradigm (adapted from Yewbrey et al., (2023), see Methods for details), from which data were analyzed for the current study. Participants were presented with a cue that signaled the preparation and production of a pre-learned sequence of 5 finger presses. An image of a black hand signaled the “Go” cue, the period in which participants could produce these sequences. Importantly, these sequences are only correct if the specific motor actions (finger presses) are conducted in a particular order and timing (like morse code or playing a piano melody), which separates these sequences in nature from abstract sequences. **C.** The abstract sequence task paradigm (adapted from Desrochers et al., (2015), see Methods for details), from which data were analyzed for the present study. At the start of each block, participants were presented with a four-item sequence (e.g., “COLOR, COLOR, SHAPE, SHAPE”), which they used to inform categorical choices about the color and shape of each stimulus. Participants kept track of each step of the rule as it occurred in time to make categorical choices, thus rendering the rules sequential in nature. Sequences were defined as abstract because they were not bound by timing or particular stimuli and required participants to impose a series of operations to respond at each trial (e.g., “COLOR, COLOR, SHAPE, SHAPE” might result in responses of “red, blue, circle, circle” on four consecutive trials) (Desrochers et al., 2022). Therefore, the identity and actions of each trial choice were not necessarily linked to the abstract sequence. At each trial, participants used their right index or middle finger to indicate their choice. Importantly, these sequences are not pre-learned, nor can participants plan their motor actions prior to each trial, which differentiates these sequences from those in the motor sequence task.

Abstract sequences, however, are supported by prefrontal neural dynamics that may generalize as a tracking mechanism across sequence types. Abstract sequences are not defined by the content of their motor actions; rather, their execution relies on following a structure or rule, such as following a recipe to bake a cake (**Figure 1A**). Previous work showed abstract sequences were supported by ramping, defined as a monotonic increase in neural activity from the beginning to the end of each sequence, in the RLPFC and other regions of the frontoparietal network (FPN) (Desrochers et al., 2015). Further, this study showed the necessity of this region for sequence completion using transcranial magnetic stimulation. The RLPFC is therefore crucial for the completion of abstract sequences. However, it remained unknown if ramping in this region supports motor sequence completion, as well, which could demonstrate that these dynamics act as a common supporting mechanism for both sequence types.

We tested the hypothesis that RLPFC ramping supports motor sequence production by analyzing data from a previously published study investigating sequences of pre-learned finger presses (**Figure 1B**) (Yewbrey et al., 2023) in comparison to a previous abstract task sequence study (**Figure 1C**) (Desrochers et al., 2015). Contrary to our hypothesis, we did not observe significant ramping activation in RLPFC during motor sequence production. However, we did observe significant ramping activation in parietal and cortical regions such as the ventromedial PFC and inferior parietal cortex, that were distinct from areas of activation from an abstract sequence task dataset (Desrochers et al., 2015). Further, we determined there was negative ramping in the frontoparietal network (FPN) during motor sequence execution compared to positive ramping in this network during abstract sequence production. Our findings suggest potentially differentiable parietal regions and network contributions underlying motor and abstract sequence production.

## Methods

In the motor sequence task, 24 participants completed the original experiment and thus were used as subjects in the present study. Details of study participants, apparatus, and data acquisition can be found in the original study (Yewbrey et al., 2023). In the abstract sequence task, 28 participants completed the original study and thus were used as subjects in the present investigation. Details of these participants and data acquisition can be found in the original study (Desrochers et al., 2015). Both tasks will be briefly described below.

### Task Design and Procedure

*Motor Sequence Task:* Participants in the original study were trained to produce four 5-finger sequences from memory in a delayed sequence production paradigm (Yewbrey et al., (2023); summarized in **Figure 1B**). Sequences had defined inter-press intervals (IPIs) and two different orders and timings, making four possible conditions. The timing structures were the same across participants (T1: 1200 ms-810 ms-350 ms-650 ms; T2: 350 ms-1200 ms-650 ms-810 ms). The experiment consisted of “Go” and “No-Go” trials. Go trials began with a sequence cue (a specific fractal image), which was associated with one of the four sequences. The cue was then followed by a screen with a fixation cross of jittered timing, which was followed by a green screen with the image of a black hand. This image cued the onset of sequence production. After the production window, another fixation screen of jittered timing appeared, followed by feedback. No-Go trials followed the same structure as Go trials but lacked the “go” cue (green screen with a black hand). Participants therefore had to remember four total fractal cues that each represented a different sequence (T1O1, T2O1, T1O2, T2O2). During the production period in No-Go trials, the fixation screen was shown for an additional 1000 ms. In the present study, we analyzed only the Go trials. A full description of the task design and procedure can be found in the original neuroimaging study (Yewbrey et al., 2023).

*Abstract Sequence Task:* Participants in the original abstract sequence study were presented at each block with four-item sequences, which they used to inform categorical choices made at every trial (Desrochers et al., (2015); summarized in **Figure 1C**). Participants had to keep track of the position of the sequence they were in, in real-time, to make correct responses about the color or shape of the stimulus at every trial. We define sequences as an ordered series of operations with a beginning and an end. Sequences were considered abstract because they are not bound by exact timing or stimuli, and participants were required to recall and impose an order of operations to make the correct response at each trial (e.g., if they were given the abstract sequence “COLOR, COLOR, SHAPE, SHAPE” they might respond “blue, red, circle, circle” to four consecutive trials) (Desrochers et al., 2022). Therefore, although motor actions are not linked, operations are still linked and ordered. Behavioral evidence supports the fact that abstract sequences are treated as a unit because of increased reaction times at sequence initiation, over and above what is typically observed for task switching and repeating (Desrochers et al., 2015, 2019; Schneider & Logan, 2006; Trach et al., 2021). In this experiment, a “trial” is considered a single item in the sequence, i.e., a single stimulus presentation. On every trial, participants were presented with a stimulus of varying size, shape (circle or square), and color (red or blue). After each trial was an intertrial interval, shown as a white fixation cross on a black screen, presented with jittered timing (0.25 – 8 s). Participants had 4 seconds on each trial to make a button response using their right index or middle fingers, which mapped to response options of the color and shape of the stimulus. Each response option was one shape and color combination (e.g., index finger button maps onto both ‘blue’ and ‘circle’ and the middle finger maps onto ‘red’ and ‘square’). Participants pressed one button per trial to indicate their response. Response options were always shown on the bottom left and right of the screen. At the end of each block, participants were asked to make a button press to indicate what they think the position of the next trial in the sequence would be. Participants completed 4 blocks (each 24-27 trials) of the task per run, for 5 runs total during fMRI scanning. Participants were trained on 3 practice sequence blocks prior to scanning. A full description of the task design and procedure can be found in the original study (Desrochers et al., 2015).

### Data Acquisition

*Motor Sequence Task Study:* A Philips Ingenia Elition X 3T MRI scanner using a 32-channel head coil was used for whole-brain imaging. T2*-weighted functional data were acquired across six runs (repetition time, TR = 2.0 s; echo time, TE = 35 ms; flip angle 90°; 60 odd-even interleaved slices; 2.0 mm isotropic voxel size) in an interleaved odd-even EPI-acquisition at a multiband factor of 2. T1-MPRAGE anatomical scans were acquired at a 0.937 × 0.937 × 1 resolution, with an FOV of 240 × 240 × 175 (A-P, R-L, F-H), encoded in the anterior-posterior dimension. See Yewbrey et al., (2023) for a full description of data acquisition.

*Abstract Sequence Task Study:* A Siemens 3T Trio Tim MRI scanner with a 32-channel head coil was used for whole-brain imaging. A high-resolution T1 weighted multiecho MPRAGE anatomical image was first collected (repetition time, TR = 2.2 s; echo time, TE = 1.54, 3.36, 5.18, 7.01 ms; flip angle 7°; 144 sagittal slices, 1.2 x 1.2 x 1.2 mm). Functional images were acquired across 5 runs using a gradient-echo echo-planar sequence (TR = 2 s; TE = 28 ms; flip angle = 90°; 38 interleaved axial slices, 3 x 3 x 3 mm). After the functional runs, a T1 in-plane anatomical image was collected (TR = 350 ms; TE = 2.5 ms; flip angle = 70°; 38 interleaved transversal slices; 1.5 x 1.5 x 3 mm). See Desrochers et al., (2015) for a full description of data acquisition.

### Data Analysis

#### Preprocessing

*Motor Sequence Task Study:* In the original study, images underwent slice time correction, realignment and unwarping, and EPI images were co-registered to each subject’s T1 anatomical scan (see Yewbrey et al., (2023) for more information). These images were further preprocessed using Statistical Parametric Mapping (SPM12) in Matlab 2023a. Specifically, images were normalized to the Montreal Neurological Institute (MNI) stereotaxic template with affine regularization. smoothed using an 8mm full-width at half-maximum Gaussian kernel, and resampled using trilinear interpolation.

*Abstract Sequence Task Study:* fMRI images were preprocessed using SPM8 (http://www.fil.ion.ucl.ac.uk/spm). Data were slice-time corrected by resampling slices in time to match the first slice. Images were then motion corrected using a rigid B-spline interpolation, and then normalized to the MNI stereotaxic space using affine regularization. Images were subsequently smoothed using an 8 mm full-width at half-maximum Gaussian kernel, and resampled using trilinear interpolation.

#### FMRI Models

All general linear models were constructed using Statistical Parametric Mapping (SPM12) software with custom scripts in Matlab 2023a. Onsets and parametric regressors were convolved with the canonical hemodynamic response function (HRF). Onset regressors were additionally convolved with the first time derivative of the HRF. Errors made during sequence production were categorized as separate events and included as onset regressors with a duration of 0 s in the models. To account for variance due to translational and rotational motion (x, y, z, roll, pitch, yaw), we included six nuisance regressors.

*Motor Sequence Task:* A subject-specific fixed-effects parametric ramp model was used to estimate beta values related to regressors. This model tests for ramping activity associated with each sequence position (1-5). We constructed regressors for each sequence condition (O1T1, O2T1, O1T2, O2T2), which included a zero-duration onset for each button press during production and a parametric (numbers 1-5) for a linear increase across the five sequence positions. Onset and parametric regressors were orthogonalized and estimated hierarchically in SPM so that variance attributed to parametric regressors was above and beyond variance accounted for by stimulus onset alone. The parametric ramp model was based on previously constructed models (Desrochers et al., 2015, 2019) used to assess ramping activity related to sequences (see description below). Examples of the unconvolved, convolved, and orthogonalized onset and parametric regressors are shown in **Figure 2**. To assess the power and validity of the parametric ramp model in the present study, we computed a Variance Inflation Factor (VIF) index (Belsley et al., 1980). VIFs were obtained by first correlating onset with parametric regressors (Pearson’s correlation coefficient). The following formula was then used to calculate a VIF per condition per participant: 1/(1 – *R^2^*). VIFs were averaged across conditions, so that each participant had one average VIF score. The average VIF per participant was 0.9885 (median 0.9884, range 0.9874 – 0.9895). Since a VIF of < 5 indicates low collinearity within a model, these results show these regressors are highly uncorrelated.

**Figure 2.**
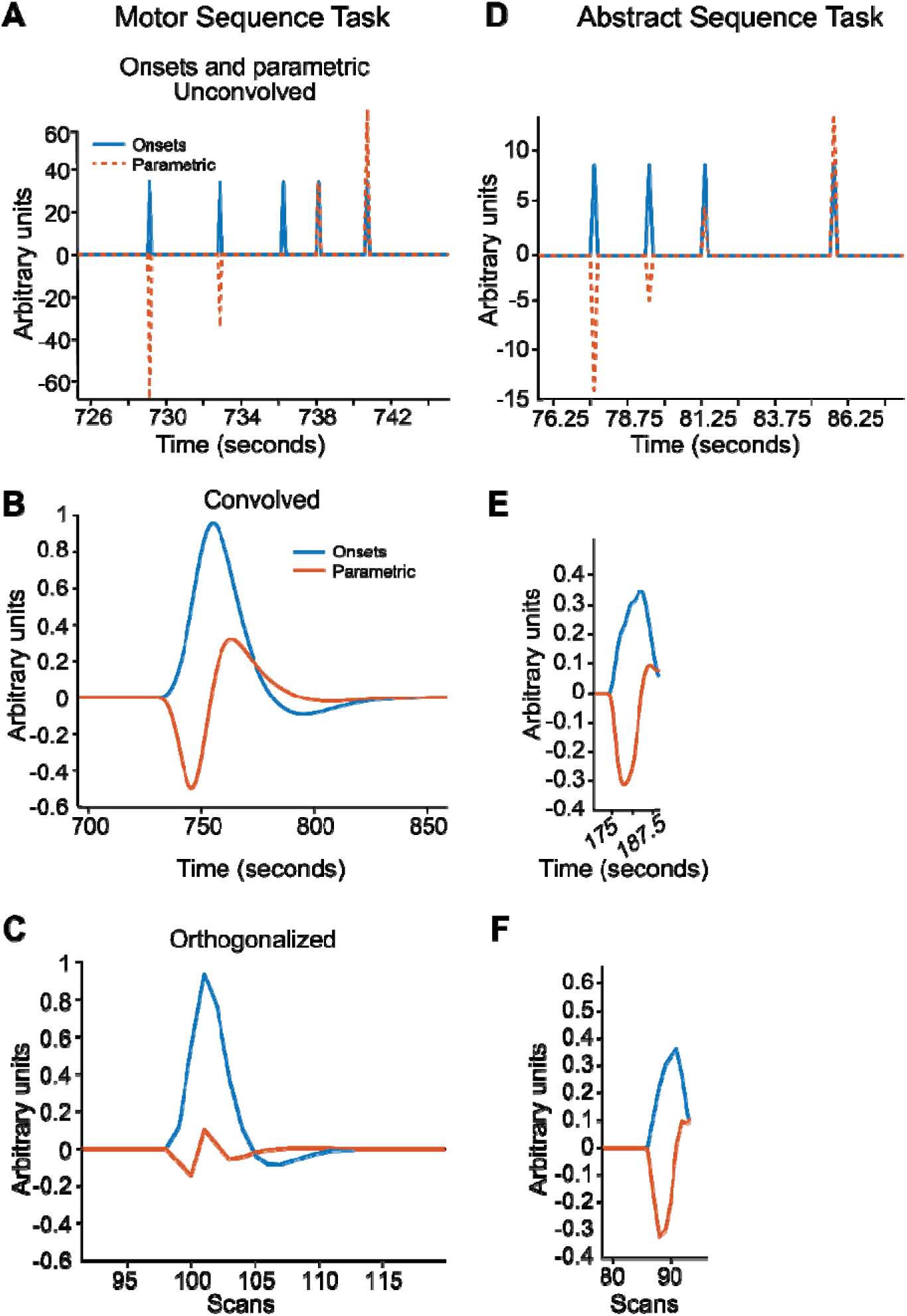
Onset and parametric regressor examples from the motor and abstract sequence studies. **A.** Example of the unconvolved onset (blue) and parametric (red) regressor for each finger press of one condition in the motor sequence task study. **B.** Example of the convolved onset and parametric regressors for one condition in the motor sequence task study. **C.** Example of the orthogonalized onset and parametric regressors for one condition of the motor sequence task study. **D.** Unconvolved onset and parametric regressors from the abstract sequence study for one sequence iteration. **E.** Convolved onset and parametric regressors for one sequence iteration from the abstract sequence study. **F.** Orthogonalized and down sampled onset and parametric regressors for one sequence iteration from the abstract sequence study. Timing in the abstract sequence study was jittered (0.25 – 8 s between stimuli), leading to differences in timing between the two tasks. However, onset and parametric regressors were orthogonalized for each task and therefore not correlated, and variance not attributed to onsets was assigned to the parametric regressors in both tasks.

*Abstract Sequence Task:* The parametric ramp model tested for ramping activity associated with each sequence position (1-4). The model was constructed as the motor sequence parametric ramp model, such that it included a zero-duration onset for each position and a parametric (number 1-4) for a linear increase across the four positions. Onset and parametric regressors were orthogonalized and estimated hierarchically in SPM, so that variance attributed to parametric regressors was above and beyond variance accounted for by stimulus onset alone. Examples of regressors from this model are shown in **Figure 2A-C**. We also calculated VIF indices to assess the validity and power of the parametric model for the abstract sequence task. The average VIF per participant was 0.9483 (median 0.9493, range 0.945 – 0.9503). These VIF scores indicate low collinearity of regressors within the model.

Although the timing between the motor and abstract sequence tasks differed, the parametric models built for both tasks are similar. The abstract sequence task contained jittered timing between trials (0.25 – 8 s, mean 2 s between stimuli). The motor sequence task had jitter as well, but comparatively less. The motor sequence task was jittered by target press timing and natural motor jitter. Participants learned sequences with one of two inter-press interval timings: either 1200 ms-810 ms-350 ms-650 ms or T2: 350 ms-1200 ms-650 ms-810 ms between finger presses. These timing differences are reflected in **Figure 2**. Despite these differences, we were able to separate the onset and parametric regressor signals from both tasks during the sequence production period (**Figure 2D-F**) to illustrate the similarities between them. Further, in both the abstract and motor sequence tasks, onset and parametric regressors were orthogonalized, such that these regressors were not correlated as indicated by the VIF scores calculated for each model, providing further evidence for the similarity of these models across both tasks.

Whole brain contrasts estimated subject-specific effects, and these estimates were then entered into a second-level analysis with subject treated as a random effect. Beta values were used for all analyses. Whole brain group voxel-wise effects were corrected for multiple comparisons using extent thresholds at the cluster level to yield family-wise error correction and were considered significant at P < 0.05. Group level contrasts were rendered on a 3D brain using Connectome Workbench (humanconnectome.org/software/connectome-workbench).

#### ROI Analysis

Regions of interest (ROIs) were constructed from previous studies, taken from whole brain analyses, or were derived from a functional network parcellation. The RLPFC (RLPFC-D15) ROI was taken from a previous abstract sequence study (Desrochers et al., 2015), defined from significant peaks of activation from the Parametric ramp model voxelwise contrast All Ramp > Baseline (referred to as Abstract Sequence Production Ramp > Baseline in the present study). The AAL atlas (Tzourio-Mazoyer et al., 2002) was used to define anatomical ROIs for the parietal cortex. We specifically used the angular gyrus and inferior parietal cortex ROIs (referred to as the AAL angular gyrus and the AAL inferior parietal ROIs) to test ramping activation during motor and abstract sequences. The peak activation-derived inferior parietal ROI (referred to as AG-RAMP ROI) was taken from whole brain analyses, derived from a significant peak of activation from the Parametric ramp model voxelwise contrast Production Ramp > Baseline in the present study. This biased, contrast-derived ROI (AG-RAMP) was used for the purpose of directly comparing ramping dynamics during motor and abstract sequence production. Lastly, the network ROIs were taken from a functional network parcellation (Yeo et al., 2011). We extracted beta values from these ROIs, and repeated measures analysis of variance (RM-ANOVA) or t-tests were subsequently performed on these values. Post-hoc analyses were conducted on RM-ANOVAs, correcting multiple comparisons using the Bonferroni method (Dunn, 1961). Effect sizes were reported as Cohen’s d (d) (Cohen, 1988) for one and two sample t-tests and partial eta squared (*η^2^*) (Cohen, 1973) for RM-ANOVAs.

## Results

In the motor sequence task, participants performed four different pre-trained motor sequences, each of five finger presses, while undergoing fMRI scanning (Yewbrey et al., 2023). Each sequence had a different inter-press timing interval and finger order. Data in the current study were analyzed from the production phase of “Go” trials when participants performed the five finger press sequences (**Figure 1B**). Each sequence was indicated at the trial onset by a specific fractal. The cue was followed by a preparation period indicated by a white fixation cross on a black screen, and then a green screen with a centered black hand icon, which indicated to the participants to initiate the finger presses (the “go” cue). The production period of each trial was followed by a fixation period and feedback on the current trial. Participants were trained prior to the fMRI scanning session to perform the four sequences from memory when cued.

In the abstract sequence task, participants were presented in each block with four-item sequences, which they used to inform categorical choices made in every trial, e.g., Color, Color, Shape, Shape, (Desrochers et al., (2015); **Figure 1C**). On every trial, participants were presented with a stimulus of varying size, shape (circle or square), and color (red or blue). Participants had 4 seconds on each trial to make a single button response to categorize the stimulus according to the position in the sequence. For example, on the first trial illustrated in **Figure 1C**, the participant is performing a “Color” categorization and therefore presses the “2” button to indicate “blue”. Response mappings remained on the screen throughout the trials. In this task, the motor response (button press) cannot be prepared ahead of time because only the categorization task is known ahead of time (according to the sequence), and not the identity of the stimulus itself. After each trial was an intertrial interval, shown as a white fixation cross on a black screen, presented with jittered timing (0.25 – 8 s). Participants completed 4 blocks (each 24-27 trials) of the task per run, for 5 runs total during fMRI scanning. Participants were trained on 3 practice sequence blocks prior to scanning.

We first tested the hypothesis that significant ramping occurs in the RLPFC during motor sequence production. We modeled a parametric ramp across finger presses for each sequence to estimate ramping activity during sequence production. This model consisted of a zero-onset duration for each button press followed by a parametric (numbers 1-5) increase across each sequence position. The onset and parametric regressors were convolved and orthogonalized, so that they were not correlated with each other (see Methods). Onset and parametric regressors were estimated in a stepwise fashion, such that variance explained by the parametric ramping regressor is above and beyond variance explained by the onsets alone. To assess ramping in RLPFC, we defined an ROI from a peak cluster of ramping activity in a previous study of healthy controls (Desrochers et al., 2015), hereafter referred to as the RLPFC-D15 ROI. Averaging across conditions, in the Motor Sequence Production Ramp > Baseline contrast we did not observe ramping activity significantly different from zero in the RLPFC-D15 ROI (**Figure 3A**; t(23) = −1.29, p = 0.21, d = −0.27). Therefore, we did not find evidence of RLPFC ramping activity during motor sequence production.

**Figure 3.**
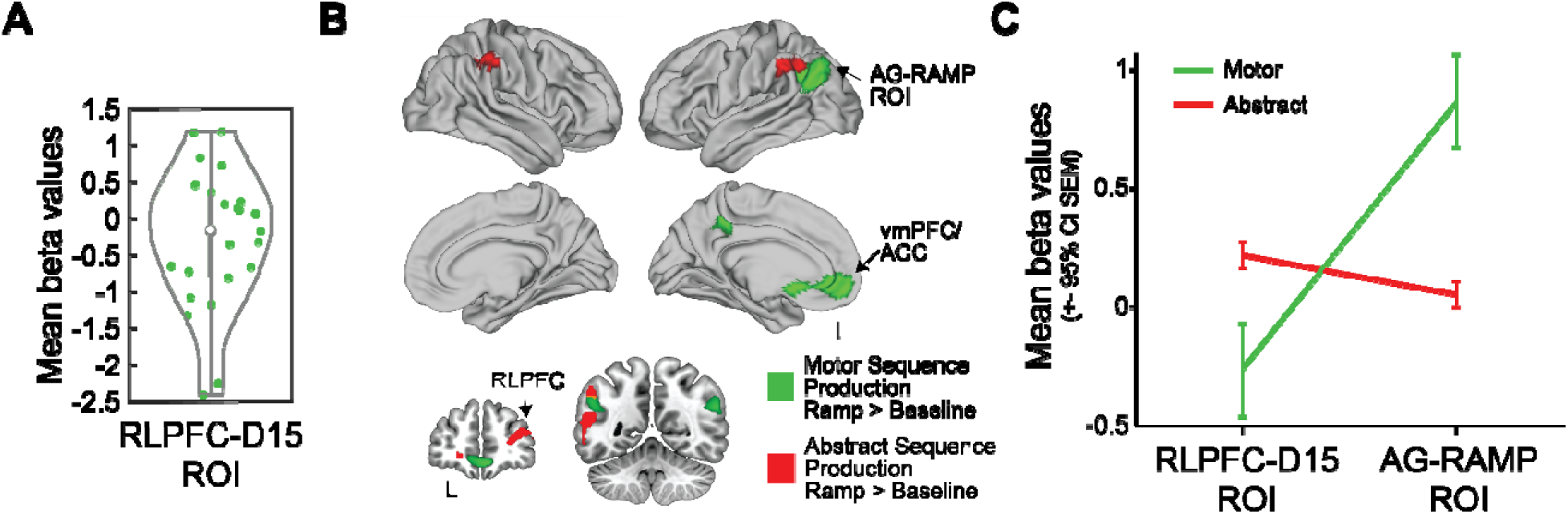
Ramping in RLPFC (RLPFC-D15) and throughout the brain during motor sequence and abstract sequence production. **A.** Average ramping activity during motor sequence production is not significantly different from zero in the RLPFC-D15 ROI. Data plotted in this graph is the same as mean ramping betas plotted for motor sequence production in the RLPFC-D15 ROI in (C) **B.** Significant ramping activity in the Abstract Sequence Production Ramp > Baseline contrast from Desrochers et al., (2015) shown in red, significant ramping activity in the Motor Sequence Production Ramp > Baseline contrast in the present study shown in green. Activity shown on whole brain renderings and in slices (FWE cluster corrected at *P < 0.05*, extent threshold 217 voxels for the Motor Sequence Production Ramp > Baseline contrast and 172 voxels for the Abstract Sequence Production Ramp > Baseline contrast. **C.** Average ramping activity in the RLPFC-D15 ROI is not significantly greater than zero during motor sequence execution but is significantly greater than zero during abstract sequence execution. Datapoint for RLPFC-D15 ROI in motor sequence production (green) is the same data as plotted in (A), reproduced here for ease of comparison. Abstract sequence ramping is significantly greater than motor sequence ramping in the RLPFC-D15. Average ramping activity in the AG-RAMP ROI is not significantly different from zero during abstract sequence execution but is significantly greater than zero during motor sequence production. Motor sequence ramping is significantly greater than abstract sequence ramping in the AG-RAMP ROI.

Since ramping during abstract sequences occurs in a network of regions beyond the RLPFC (Desrochers et al., 2015), we next tested if significant ramping occurs in other regions of the whole brain during motor sequence production. We examined the whole brain Motor Sequence Production Ramp > Baseline contrast and observed three significant clusters in ventromedial PFC (vmPFC) and in the inferior parietal cortex (**Figure 3B**; **Table 1**). We compared these clusters with regions of ramping observed during abstract sequences and found that these clusters did not overlap with ramping activity previously observed in the whole brain Abstract Sequence Production Ramp > Baseline contrast during abstract sequence execution (Desrochers et al., 2015). From these contrasts, we showed significant ramping occurred during motor sequence production in different regions than ramping during abstract sequences.

**Table 1.**
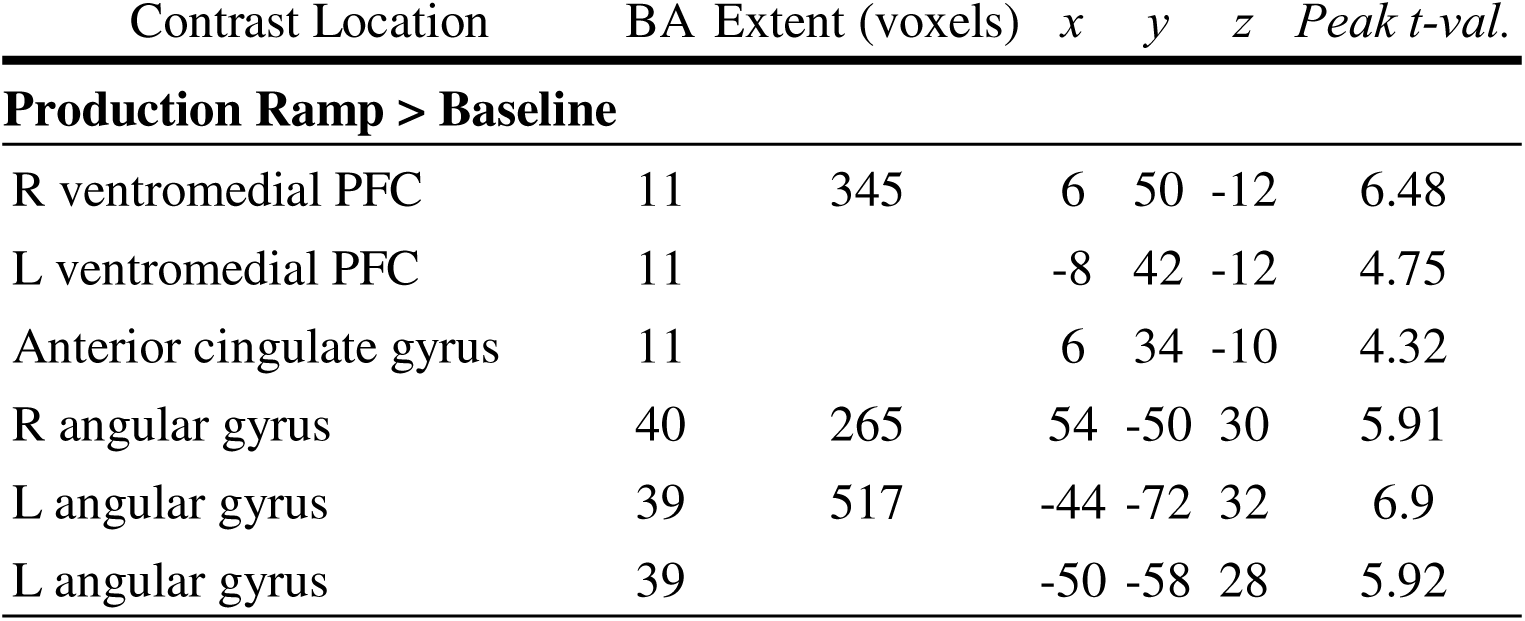
Activation coordinates, significant neural activity in the Production Ramp > Baseline contrast. Clusters reliable at p < 0.05 corrected. Coordinates are the center of mass in MNI. Clusters are reported for peaks of activation 8 mm or greater distance apart.

| Contrast Location | BA | Extent (voxels) | *x* | *y* | *z* | *Peak t-val.* |
| --- | --- | --- | --- | --- | --- | --- |
| **Production Ramp > Baseline** |  |  |  |  |  |  |
| R ventromedial PFC | 11 | 345 | 6 | 50 | -12 | 6.48 |
| L ventromedial PFC | 11 |  | -8 | 42 | -12 | 4.75 |
| Anterior cingulate gyrus | 11 |  | 6 | 34 | -10 | 4.32 |
| R angular gyrus | 40 | 265 | 54 | -50 | 30 | 5.91 |
| L angular gyrus | 39 | 517 | -44 | -72 | 32 | 6.9 |
| L angular gyrus | 39 |  | -50 | -58 | 28 | 5.92 |

The bilateral region of inferior parietal cortex ramping during motor sequence production appears to largely lie in the angular gyrus. Therefore, in the remaining analyses, this activation cluster will be referred to as the AG-RAMP ROI (as labeled in **Figure 3B,C**).

To more directly test ramping activity differences between motor sequence and abstract sequence production, we compared ramping from both datasets in the RLPFC-D15 and the AG-RAMP ROIs. Overall ramping beta values were greater in the AG-RAMP than in the RLPFC-D15 (ROI main effect: F(1,49) = 23.57, p < 0.001, η^2^ = 0.32) and effects in each ROI were significantly different across the two sequence tasks (ROI*sequence type interaction: F(1,49) = 43.56, p < 0.001, η^2^ = 0.47). Follow up post-hoc tests showed that ramping beta values during motor sequences were not significantly different from zero in the RLPFC-D15 (t(23) = −1.38, p = 0.36, d = −0.27; **Figure 3C** in green). Abstract sequence ramping was significantly greater than zero in the RLPFC-D15 ROI (t(27) = 4.43, p < 0.001, d = 0.83; **Figure 3C** in red), and significantly greater than motor sequence ramping in this ROI (t(49) = 2.62, p = 0.02, d = 0.72; **Figure 3C**, left side). We further observed that abstract sequence ramping was not significantly greater than zero in the AG-RAMP ROI (t(27) = 1.66, p = 0.22, d = 0.06; **Figure 3C**, in red), and that motor sequence ramping was significantly greater than zero (t(23) = 6.24, p < 0.001, d = 0.87; **Figure 3C** in green) and significantly greater than abstract sequence ramping in this ROI (t(49) = −6.08, p < 0.001, d = −1.68; **Figure 3C**, right side). These results show differentiable ramping patterns of the prefrontal and parietal cortex between sequence types, such that RLPFC (RLPFC-D15) ramping is specific to abstract sequence execution while ramping in the angular gyrus (AG-RAMP ROI) occurs during motor and not abstract sequence execution.

To more directly test if regions of parietal ramping during motor sequences are distinct from those that ramp during abstract sequences, we conducted further analyses in parietal subregions using anatomical ROIs. We used the angular gyrus and inferior parietal ROIs, defined by the AAL atlas (Tzourio-Mazoyer et al., 2002) (**Figure 4A**). We analyzed the left and right hemispheres separately since parietal ramping activation in both sequence tasks did not occur in the same regions bilaterally.

**Figure 4.**
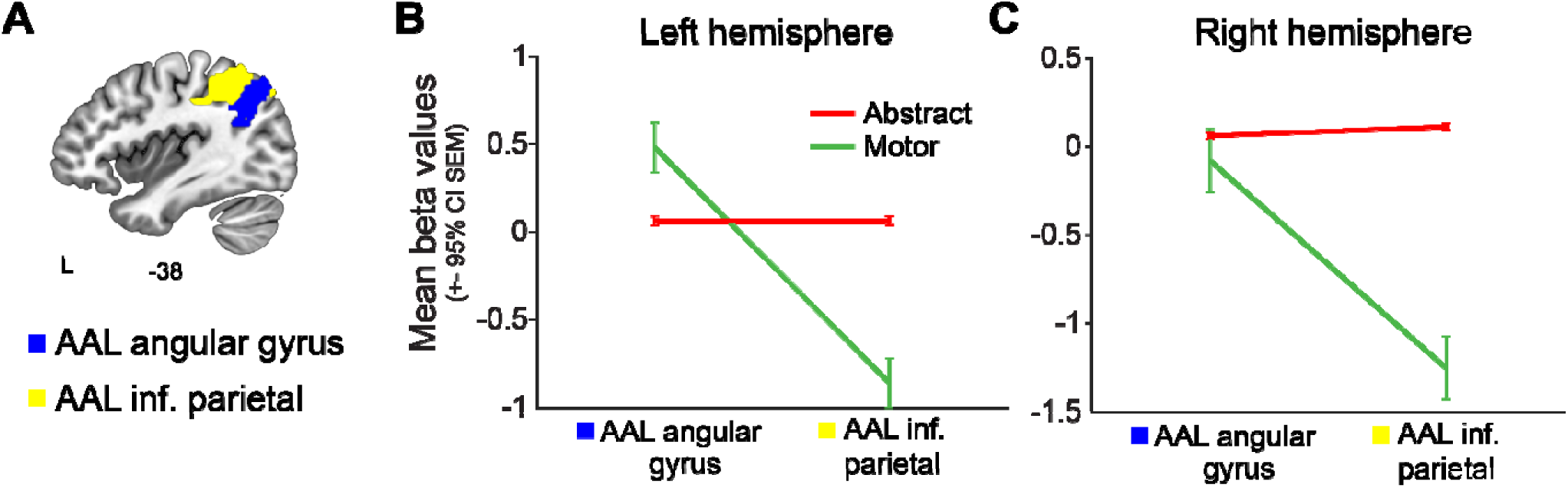
Ramping during motor and abstract sequence production in anatomical atlas-based ROIs of the inferior parietal cortex. **A**. The anatomical AAL ROIs used for analysis were the angular gyrus (blue) and inferior parietal (yellow), illustrated for example on a left hemispheric sagittal slice. These ROIs were chosen as they largely cover the ramping activation clusters observed during abstract and motor sequence execution but are unbiased anatomical delineations of the inferior parietal cortex. **B.** Average beta values from the Abstract Sequence Production > Baseline (red) and the Motor Sequence Production > Baseline (green) in the left AAL angular gyrus and inferior parietal ROIs. There was a significant interaction such that there was increased ramping in motor compared to abstract in the angular gyrus but increased ramping in abstract compared to motor in the inferior parietal ROI. **C.** Average beta values from the Abstract Sequence Production > Baseline (red) and the Motor Sequence Production > Baseline (green) in the right AAL angular gyrus and inferior parietal ROIs. There was a significant interaction between sequence types by each ROI, driven by increased ramping in the right AAL inferior parietal during abstract compared to motor sequence execution.

Overall, in the left hemisphere, ramping differentiated the two sequence types between the two AAL ROIs (**Figure 4B**). There was a significant difference in ramping beta values between motor and abstract sequences (Sequence type main effect: F(1,49) = 5.31, p = 0.03, η^2^ = 0.10) and between the AAL angular gyrus and AAL inferior parietal ROIs (ROI main effect: F(1,49) = 92.77, p < 0.001, η^2^ = 0.65). Furthermore, ramping significantly differed by sequence type between each ROI (Sequence type*ROI interaction: F(1,49) = 93.4, p < 0.001, η^2^ = 0.66). To further examine the details of this interaction, we performed post-hoc analyses. Within the left AAL angular gyrus ROI, ramping was significantly greater than zero during motor sequence execution (t(23) = 3.92, p < 0.001, d = 0.77), but activity during abstract sequences was not significantly greater than zero (t(27) = 1.93, p = 0.12, d = 0.36). Within the left AAL inferior parietal ROI, motor sequence ramping was significantly less than zero (t(23) = −5.96, p < 0.001, d = −1.18). Negative ramping, a ramping signal significantly less than zero, indicates that there is activity at the start of the motor sequence that significantly decreases across the production period. Abstract ramping in the left AAL inferior parietal ROI was significantly greater than zero (t(27) = 2.54, p = 0.03, d = 0.47). Between the two tasks, motor sequence ramping was significantly greater than abstract sequence ramping in the left AAL angular gyrus ROI (t(49) = −3.48, p = 0.002, d = −0.96), while abstract sequence ramping was significantly greater than motor sequence ramping in the left AAL inferior parietal ROI (t(49) = 6.68, p < 0.001, d = 1.84; **Figure 4B**). This interaction supports differentiable ramping dynamics between motor and abstract sequences in the AAL angular gyrus and AAL inferior parietal cortex ROIs in the parietal cortex.

A similar interaction in the right hemisphere supports differentiation between sequence types as well, although not as strongly as in the left hemisphere. Overall, in the right hemisphere, ramping beta values were significantly different between sequence types (Sequence type main effect: F(1,49) = 27.33, p < 0.001, η^2^ = 0.36) and of ROI (F(1,49) = 42.41, p < 0.001, η^2^ = 0.46). Furthermore, there was a significant difference in ramping between sequence types across both ROIs (Sequence type*ROI interaction: F(1,49) = 50.45, p < 0.001, η^2^ = 0.51). To further examine the details of this interaction, we performed post-hoc analyses. Within the right AAL angular gyrus, motor sequence ramping was not significantly different from zero (t(23) = −0.49, p = 0.63, d = −0.10), while abstract sequence ramping was significantly greater than zero (t(27) = 2.44, p = 0.04, d = 0.46). Within the right AAL inferior parietal ROI, motor sequence ramping was significantly less than zero (t(23) = −6.63, p < 0.001, d = −1.31), while abstract sequence ramping was significantly greater than zero (t(27) = 3.77, p < 0.001, d = 0.71). Between the two tasks, ramping was not significantly different in the right AAL angular gyrus (t(49) = 0.91, p = 0.73, d = 0.25), but abstract sequence ramping was significantly greater than motor sequence ramping in the right AAL inferior parietal ROI (t(49) = 7.55, p < 0.001, d = 2.09; **Figure 4C**). This interaction therefore supports a partial differentiation of sequence types by the two anatomical parietal ROIs in the right hemisphere.

Given the potential differences in ramping across hemispheres, we compared the left and right hemispheres to determine if there was a difference in ramping between them. We found that overall, ramping activity in the left hemisphere was greater than ramping activity in the right hemisphere (main effect hemisphere: F(1,99) = 5.12, p = 0.03, η^2^ = 0.05) with no interaction by ROI (hemisphere*ROI: F(1,99) = 0.89, p = 0.34, η^2^ = 0.01). Therefore, although the overall level of ramping activity is different across hemispheres, these results suggest that the pattern of ramping activity and crossover interactions between the ROIs are not different across hemispheres. We therefore provide evidence using unbiased anatomical ROIs that different regions of the parietal cortex, the angular gyrus and the inferior parietal cortex, are recruited via ramping dynamics during motor and abstract sequence execution in these tasks.

To test if specific networks are uniquely involved in each task, we examined motor and abstract sequence ramping in previously defined cortical networks. This investigation was motivated in part by the previous finding that there is significantly increased ramping in regions that lie largely the frontoparietal network (FPN) during abstract sequence production (Desrochers et al., 2015). To test this question, we examined ramping activity in both the abstract sequence task and motor sequence task datasets in seven previously defined cortical networks (Yeo et al., 2011) (**Figure 5A**). Ramping beta values were significantly greater across networks in abstract compared to motor sequences (Sequence type main effect: F(1,49) = 17.57, p < 0.001, η^2^ = 0.26), and ramping was significantly different across networks (Main effect of network: F(1,49) = 24.41, p < 0.001, η^2^ = 0.33). Furthermore, effects in each network were significantly different across each sequence type (Network*Sequence type interaction: F(1,49) = 25.01, p < 0.001, η^2^ = 0.33). To further investigate this interaction, we conducted post-hoc analyses to examine ramping activity in networks within each sequence type and between sequence types. During abstract sequence execution, there was significantly increased ramping in the FPN (t(27) = 3.36, p = 0.005, d = 0.62), and no significant ramping (i.e., beta values not significantly different than zero) in any other network (**Figure 5B**). During motor sequence execution, significant ramping was negative, indicating a monotonic decrease in signal from the start to the end of the sequence. Specifically, significant negative ramping (compared to zero) was located in the visual (t(23) = −5.29, p <0.001, d = −1.04), somatomotor (t(23) = −4.79, p < 0.001, d = −0.94), dorsal attention (t(23) = −7.99, p < 0.001, d = −1.58), ventral attention (t(23) = −2.96, p = 0.01, d = −0.58), and frontoparietal (t(23) = −4.15, p < 0.001, d = −0.82) networks (**Figure 5C**). Notably, ramping in the FPN was significantly greater during abstract compared to motor sequence production (t(50) = −4.25, p < 0.001, d = 1.38). We therefore show potentially opposing network dynamics in the FPN during abstract compared to motor sequence production, suggesting differential network contributions to the two sequence types during the two different tasks.

**Figure 5.**
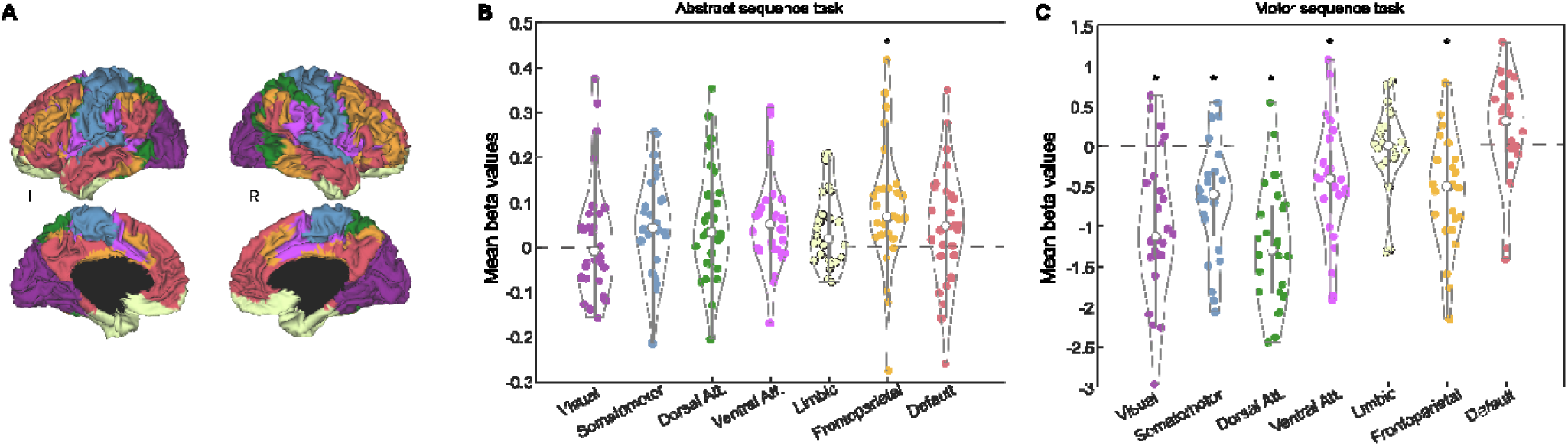
Opposing network dynamics in the frontoparietal network during abstract vs. motor sequence production. **A.** The 7 resting state cortical networks (Yeo et al., 2011) represented on 3D left and right brain hemispheres. **B.** Significant positive ramping in the FPN during abstract sequence production. **C.** Significant negative ramping in the visual, somatomotor, dorsal attention, ventral attention, and frontoparietal network during motor sequence production. FPN ramping is significantly greater during abstract compared to motor sequence production.

## Discussion

We investigated ramping dynamics during motor and abstract sequence production, using data from a previous motor (Yewbrey et al., 2023) and an abstract (Desrochers et al., 2015) sequence study, with a hypothesis that RLPFC ramping dynamics generalize to track motor in addition to abstract sequence production. Contrary to this hypothesis, we did not observe significant RLPFC ramping during motor sequences, suggesting RLPFC dynamics are specific to abstract sequences. However, we found significant motor sequence ramping in the inferior parietal cortex (AG-RAMP ROI), an area distinct from parietal ramping that occurs during abstract sequence production. Further, we observed negative FPN ramping activity during motor sequence production, in contrast to positive FPN ramping observed during abstract sequences. Overall, ramping dynamics reveal that distinct prefrontal-parietal regions and differential cortical network dynamics may support motor compared to abstract sequence production.

Since RLPFC ramping does not significantly support motor sequence production, recruitment of this region may be specific to non-motor, such as abstract, sequences. The anterior PFC has been previously associated with hierarchical abstraction and structures (Koechlin et al., 1999, 2003; Sakai and Passingham, 2003; M et al., 2005). One theory poses a hierarchical organization of the PFC (Badre and D’Esposito, 2007), such that rostral PFC regions are recruited for more abstract facets of cognitive control. Regions caudal to PFC regions such as the motor and premotor cortex, however, are recruited during motor sequence execution (Tanji and Shima, 1994; Yokoi and Diedrichsen, 2019; Russo et al., 2020). Since the motor sequences in the present study are pre-trained and memorized prior to the fMRI scan, monitoring their execution may not be required in the same manner as for abstract sequences. Therefore, in the present behavioral paradigm, rostral PFC regions such as the RLFPC may not be necessary to help monitor the production of such sequences. More generally, RLPFC may not be recruited during skilled motor sequence production. Overall, our work provides evidence for circuitry supporting pre-trained, skilled motor sequence production that is distinct from that underlying abstract sequences.

The AG-RAMP ROI and FPN ramping findings suggest differential cortical network dynamics underlying motor sequence compared to abstract sequence production. Prior neuroimaging experiments have provided some evidence for the hierarchical differentiation of parietal cortex. For example, the dorsal region of the abstract sequence parietal ramping was shown previously to hold information about both higher and lower level contextual information needed to hold memory during a task (Nee and Brown, 2012). Further, this dorsal parietal region is part of the FPN and is active during task-switching (Periáñez et al., 2024), which is necessary for flexible behavior related to higher-level cognitive processing. The AG-RAMP ROI is involved in more concrete aspects of sequential processing, such as number processing (Göbel et al., 2001; Grabner et al., 2009). These experiments suggest that parietal regions involved in abstract versus motor sequences may be differentiated by the hierarchical information they represent. In the current study, differential network dynamics, significant positive FPN ramping during abstract and significant negative FPN ramping during motor sequences, suggests opposing network dynamics may differentiate the two sequence types as well. Negative ramping, the decrease in activity across the sequence, may functionally signify the initiation of control or motor-related cortical regions needed to start the motor sequence that dissipates over the course of producing an already rehearsed repertoire. Previous work to support this idea shows that the pre-SMA is involved in initiating overly trained motor sequences (Kennerley et al., 2004). Alternatively, negative ramping could be due to decreasing activity across time in response to similar motor actions (i.e., repetition suppression between finger presses) (de C. Hamilton and Grafton, 2008; Berlot et al., 2021). Distinct parietal regions and differential cortical network dynamics may therefore support motor compared to abstract sequence production.

We observed ramping in vmPFC during motor sequences, which may be due to feedback during the task. Ramping activity occurs in anticipation of reward in the vmPFC (Lamichhane et al., 2022). During the motor sequence task, participants were provided with feedback after each production period, which may produce ramping signals similar to those seen in reward anticipation. Therefore, it is possible that vmPFC ramping observed may be more related to feedback than to sequential information during the production period. The abstract sequence task did not provide feedback to participants (Desrochers et al., 2015), which may account for the lack of ramping activity observed in the vmPFC. Future studies should be designed to isolate reward as a variable to more accurately assess ramping related directly to motor sequence production.

Limitations to this study largely involved design differences between the abstract and motor sequence task studies. As described in the Methods, the studies differed in sequence production timing, variance, and regressor correlation. Additionally, the motor sequence task contained feedback while the abstract sequence task did not. These differences may contribute to the present results, so direct comparisons between tasks and interpretation of results should be made with caution. However, despite these differences, we established low collinearity between onset and parametric regressors within each task and a low variance inflation factor (VIF, < 1) for both the motor and abstract sequence parametric ramp models (see Methods), validating the use of these models in each task. Further, we give compelling support for the separability and comparison of the onset and parametric regressors for both tasks despite timing differences (**Figure 2**). We also note that despite differences, both tasks required finger press responses and responses to each sequential step in time, which are commonalities we believe allow for the comparisons we made. Although the present study provided strong evidence for a differentiable pattern of ramping dynamics between motor and abstract sequences across two different task designs, future experiments should directly investigate ramping in motor compared to abstract sequence production using hierarchical/nested structures (potentially building on Yokoi et al., (2019) but extending to non-motor sequences). Lastly, we were unable to model the sequence preparation period parametrically without button press timings, so the present study focused only on periods of motor sequence production. Future studies should directly test ramping during motor sequence preparation compared to production. Future work may also prioritize investigating the connectivity of the differentiated parietal cortical regions in abstract versus motor sequences to better determine networks that contribute to the production of each sequence type.

Our work has implications for psychiatric disorders associated with sequential deficits, such as obsessive-compulsive disorder (OCD). Previous studies showed reaction time deficits in OCD during implicitly learned motor sequences (Kathmann et al., 2005; Kelmendi et al., 2016; Soref et al., 2018) and dysfunctional prefrontal and basal ganglia circuitry that underlie this process (Milad and Rauch, 2012; Janacsek et al., 2020). We have shown that abstract sequence execution in an OCD population is largely similar to that of healthy controls, with no difference in RLPFC activation or ramping dynamics between groups (Doyle et al., 2024). These studies suggest that representation of abstract sequence information may be preserved in OCD, but sequential deficits at the level of individual motor actions may contribute to pathological behaviors. Our present results extend this hypothesis to include specific prefrontal and parietal regions of ramping that may be dysfunctional in OCD during sequential behavior.

Overall, motor sequence production is supported by distinct prefrontal-parietal regions and opposing network dynamics compared to that of abstract sequence execution. Our results provide evidence for novel neural circuitry underlying skilled motor sequences and provide further evidence for neural differentiation of hierarchical sequence information.

## Data Availability

Imaging data are not publicly available. Analysis code is available upon request.

## Author Contributions

Hannah Doyle: Conceptualization, Investigation, Methodology, Formal Analysis, Writing – Original Draft, Writing – Reviewing and Editing, Visualization. Rhys Yewbrey: Data Curation, Conceptualization, Writing – Reviewing and Editing. Katja Kornysheva: Conceptualization, Validation, Writing – Reviewing and Editing, Supervision. Theresa M. Desrochers: Conceptualization, Validation, Writing – Original Draft, Writing – Reviewing and Editing, Supervision, Funding Acquisition.

## Acknowledgements

We thank members of the Desrochers and Kornysheva labs for their advice and feedback on the manuscript.

